# Regulation of olfactory associative memory by the circadian clock output signal Pigment-dispersing factor (PDF)

**DOI:** 10.1101/2020.04.17.046953

**Authors:** Johanna G. Flyer-Adams, Emmanuel J. Rivera-Rodriguez, Jacob D. Mardovin, Junwei Yu, Leslie C. Griffith

## Abstract

Dissociation between the output of the circadian clock and external environmental cues is a major cause of human cognitive dysfunction. While the effects of ablation of the molecular clock on memory have been studied in many systems, little has been done to test the role of specific clock circuit output signals. To address this gap, we examined the effects of mutation of *Pigment-dispersing factor (Pdf)* and its receptor, *Pdfr* on associative memory in male and female *Drosophila*. Loss of PDF signaling significantly decreases the ability to form associative memory. Appetitive short-term memory (STM), which in wildtype is time-of-day (TOD)-independent, is decreased across the day by mutation of *Pdf* or *Pdfr*, but more substantially in the morning than in the evening. This defect is due to PDFR expression in adult neurons outside the core clock circuit and the mushroom body Kenyon cells. The acquisition of a TOD difference in mutants implies the existence of multiple oscillators that act to normalize memory formation across the day for appetitive processes. Interestingly, aversive STM requires PDF but not PDFR, suggesting that there are valence-specific pathways downstream of PDF that regulate memory formation. These data argue that the circadian clock uses circuit-specific and molecularly diverse output pathways to enhance the ability of animals to optimize responses to changing conditions.

**SIGNIFICANCE STATEMENT:** From humans to invertebrates, cognitive processes are influenced by organisms’ internal circadian clocks, the pace of which is linked to the solar cycle. Disruption of this link is increasingly common (e.g. jetlag, social jetlag disorders) and causes cognitive impairments that are costly and long-lasting. A detailed understanding of how the internal clock regulates cognition is critical for the development of therapeutic methods. Here, we show for the first time that olfactory associative memory in *Drosophila* requires signaling by Pigment-dispersing factor (PDF), a neuromodulatory signaling peptide produced only by circadian clock circuit neurons. We also find a novel role for the clock circuit in stabilizing appetitive sucrose/odor memory across the day.

## INTRODUCTION

Cognition is influenced by the circadian clock. Within individual cells, time is kept by a molecular clock; the coordination of time across tissues in the organism is performed by the ‘core clock’, a small population of individually cycling neurons bound together to form a coherent circuit. The outputs of this circuit generate rhythms of physiology and behavior that oscillate with a period aligned to the solar cycle. Circadian peaks and troughs in learning and memory have been documented in humans as well as rodent and invertebrate models (Gerstner & Yin, 2010; Wright et al., 2012) such that misalignments between an organism’s cycling internal clock and external conditions (jetlag, social jetlag, and shift work disorders) impair cognition/memory across phyla (Kott et al., 2012; Weingarten & Collop, 2013; Wittmann et al., 2006; Wright et al., 2013). Irreversible neurodegenerative diseases that affect cognition such as Parkinson’s and Alzheimer’s exhibit comorbid circadian dysfunction (Videnovic et al., 2014). Even natural aging degrades the fidelity of the body’s clock in a manner that has been linked to cognitive decline (Reinhart & Nguyen, 2019). Thus a complete understanding of how the body’s clock regulates cognition could generate new therapeutic approaches.

To date, the clock-memory link has largely been investigated by breaking the transcriptional feedback loop providing intracellular timekeeping in cells of the clock circuit. Eliminating cycling with altered light conditions affects novel object recognition in rodents (Ruby et al., 2013) and time-of-day (TOD)-dependent associative memory in *Drosophila* (Chouhan et al., 2015; Le Glou et al., 2012; Lyons & Roman, 2009). Clock gene mutations such as *Per1, Per2* and *Bmal1* generate impairments in contextual fear and spatial memory tasks in mice and are involved in all phases of memory processing (Rawashdeh et al., 2014; Snider & Obrietan, 2018; Wang et al., 2009; Wardlaw et al., 2014). In *Drosophila*, mutations of the clock genes *period* and *clock* impair TOD memory and can disrupt memory generally (Chouhan et al., 2015; Fropf et al., 2014, 2018; Le Glou et al., 2012; Lyons & Roman, 2009; Sakai et al., 2004). But studies using molecular clock mutants don’t fully replicate conditions that exist in common human clock-related cognitive disorders; in most, a functional molecular clock is present but outputs are misaligned with environmental cues. To fully understand the role of clock circuit outputs on cognition, it is necessary to manipulate output pathways in the context of an intact molecular clock.

Environmentally-cued circadian rhythms of behavior and physiology are generated by the outputs of the core clock circuit which are organized by peptidergic signaling from a few pacemaker neurons (Aton et al., 2005). In *Drosophila*, this peptide is Pigment-dispersing factor (PDF), an 18 aa peptide produced in the adult brain solely by 16 ventrolateral clock neurons (LNvs) (Park et al., 2000; Renn et al., 1999). PDF released from the LNvs signals within the core clock circuit through its only known receptor PDFR (Hyun et al., 2005; Lear et al., 2005; Mertens et al., 2005; Shafer et al., 2008) to coordinate the activity of the ∼150 core clock neurons (Liang et al., 2016; Lin et al., 2004; Peng et al., 2003; Stoleru et al., 2005; Yoshii et al., 2009). Core clock circuit output directs rhythmic aspects of physiology such as locomotor activity and sleep, and PDF also functions in these output pathways. In this way, the PDF/PDFR signaling pathway both maintains free-running circadian activity and promotes output behaviors.

Here, we investigate the involvement of PDF signaling in *Drosophila* memory, examining the role of clock output on associative olfactory memory for the first time in the context of a functional molecular clock. We find a novel adult-specific requirement for PDF/PDFR signaling in memory that is distinct from its role in synchronizing the clock circuit. We also provide evidence for the existence of a second PDF receptor that allows valence-specific regulation of associative olfactory memory by PDF.

## MATERIALS AND METHODS

### Fly Stocks and Husbandry

All experimental flies were collected directly after eclosion and maintained in incubators at 25°C with a 12:12 LD cycle. Flies were reared and housed on cornmeal dextrose food with yeast except those used for GeneSwitch experiments. For Figure 1 and 6 experiments, mixed males and females were used; for cell-specific PDFR expression memory experiments and all activity experiments, males were used; for imaging experiments, females were used. Fly strains used include: *Canton-S* wildtype (WT; Canton, Ohio), ;;*pdf*^*01*^ (BDSC#26654; backcrossed 6x into WT), *han*^*5304*^;; (BDSC#33068; backcrossed 6x into WT), ;;*nsyb-GAL4/TM6B* (Goodwin et al., 2018), *han*^*5304*^;;*UAS-pdfr-myc13* (gift of Paul Taghert), ;;*elav-GS-GAL4* (Osterwalder et al., 2001), *VT030559-GAL4* (VDRC ID# v206077), *pdfR-2A-LexA;;* and ;;*pdf*^*attP*^ (Deng et al., 2019), *w*^*-*^,*UAS-mCD8-IVS-RFP,LexAop-mCD8-IVS-GFP;;* (generated from BDSC#61681), ;*pdf-GAL4;* (Park et al., 2000), *UAS-P2X*_*2*_ (Lima & Miesenböck, 2005), *MB-LexA* (Pitman et al., 2011), ;*LexAop-EPAC;* (BDSC#76031), ;*pdfR(10kb)-GAL4*; (Parisky et al., 2016), ;*clk856-GAL4;* (gift from M. Rosbash).

**Figure 1.**
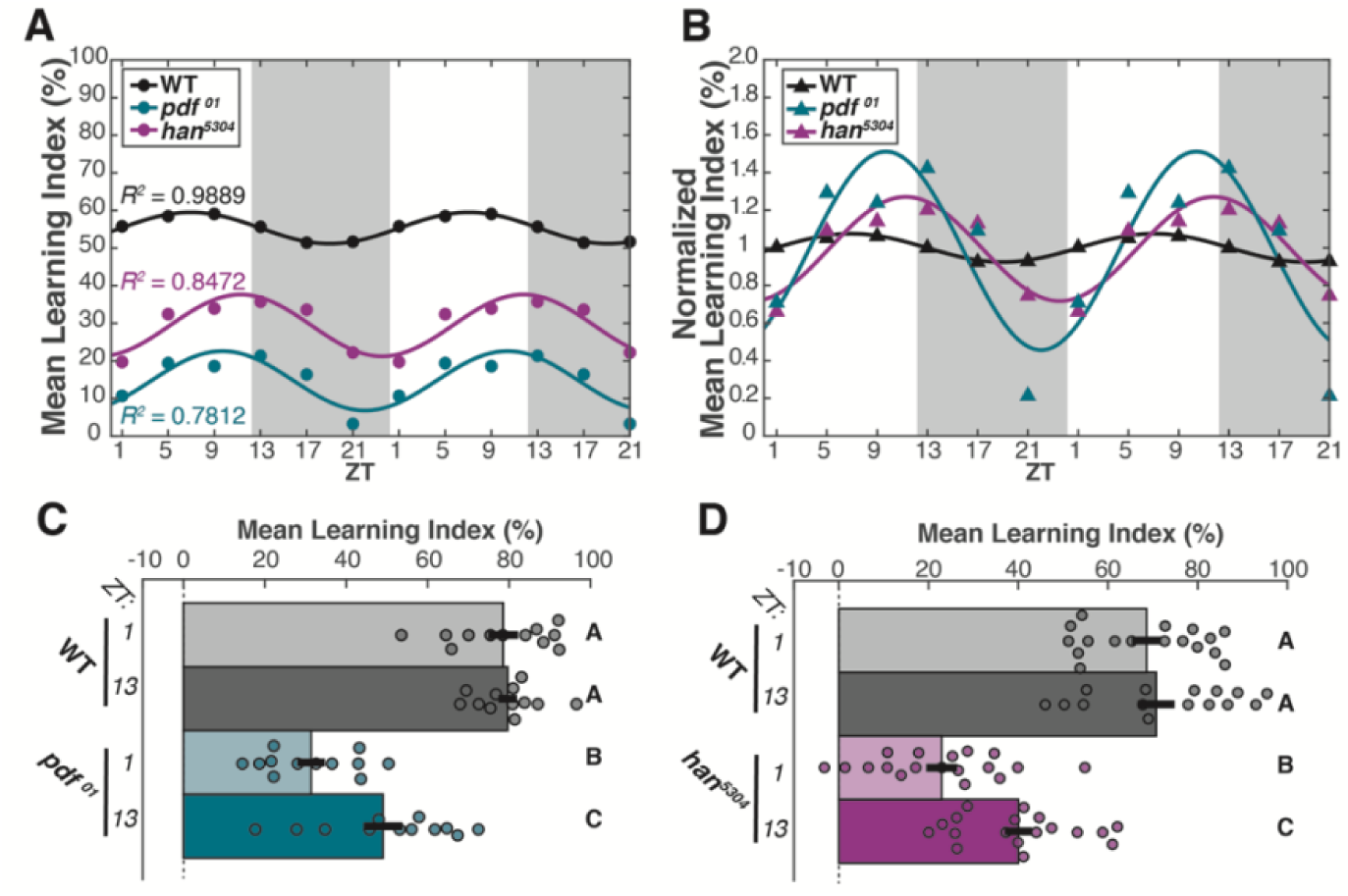
The core clock supports appetitive short-term memory throughout the circadian cycle via a PDF signaling pathway. Appetitive olfactory 2 min short-term memory (STM) of WT and PDF pathway mutants. (**A**,**B**) Learning index (LI) scores for STM tested every 4 h through the 24 h cycle and double-plotted, with white and grey background indicating lights on/off of entrainment phase. Each genotype was fit with 1-term Fourier curve, R^2^ values shown in (A). **(A)** Mean LI (circles; n=8 for each timepoint) and **(B)** normalized mean LI (triangles; mean subtracted, then divided by standard deviation). (**C**,**D**) STM of *pdf*^*01*^ and *han*^*5304*^ mutants compared to WT at ZT1 and ZT13. Mean LI scores are shown ± SEM, with individual datapoints (circles). Letter categorization indicates groups of statistical equivalence (p>0.05) or difference found by 2-way ANOVA with interactions, Bonferroni posthoc comparisons (all significant comparisons, panel C: p<0.01, panel D: p<0.005).

**Figure 2.**
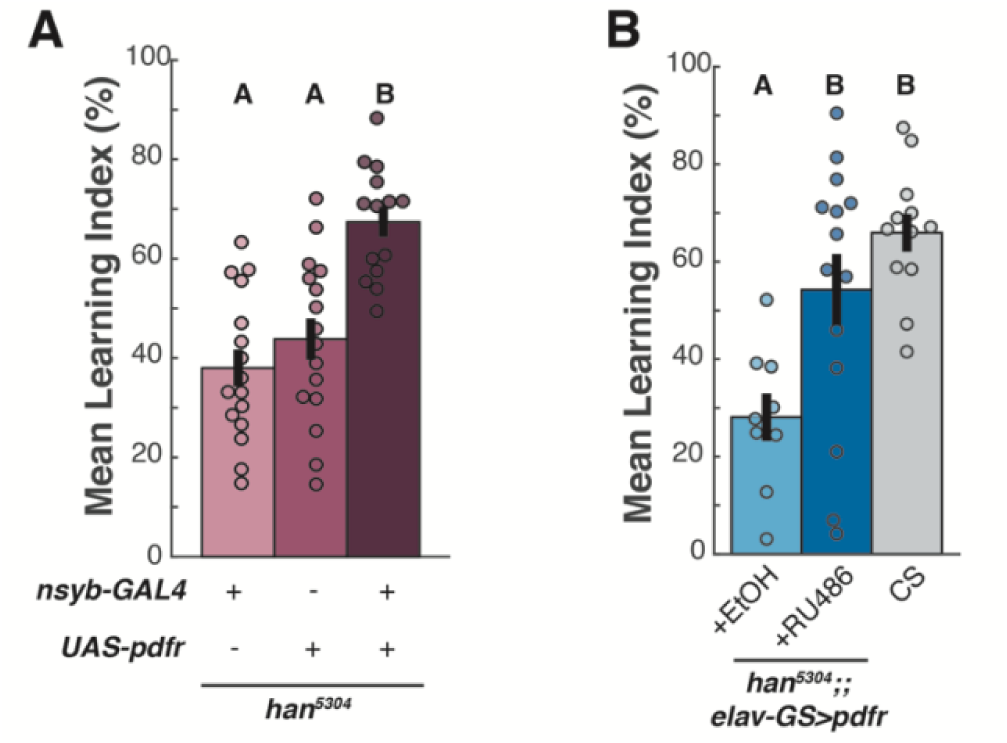
Adult-specific neuronal PDF-PDFR signaling is sufficient for appetitive short-term memory formation. Appetitive olfactory STM tested between ZT0-4 in *han*^*5304*^ mutants with panneuronal PDFR expressed (**A**) constitutively, compared to genetic controls and (**B**) adult-specifically using the RU486-inducible GeneSwitch system, compared to EtOH vehicle and WT controls. Mean LI scores are shown ± SEM, with individual datapoints (circles). Letter categorization indicates groups of statistical similarity (p>0.05) or difference found by 1-way ANOVA with Bonferroni posthoc comparisons (all significant comparisons, panel A: p<0.0005, panel B: p<0.05).

**Figure 3.**
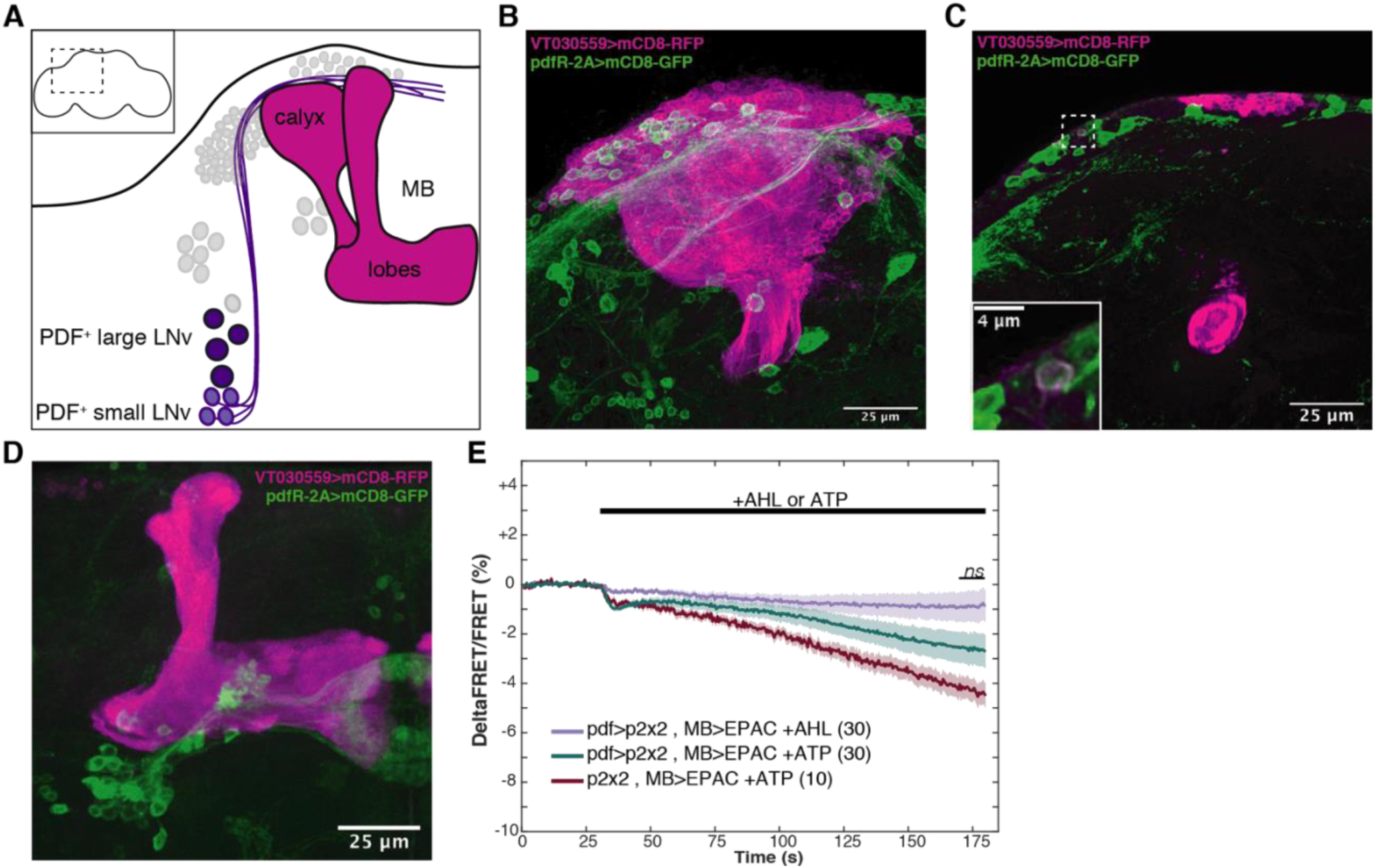
The MB is not a direct target of the PDF signaling pathway. **(A)** Cartoon representation of LNv/MB anatomy. Purple PDF^+^ LNv soma (light:small, dark:large) are shown with other core clock cells in light grey. sLNvs send projections anterodorsomedially in close proximity to the MB (magenta) calyx, as indicated. (Inset) Area shown in main panel indicated by dotted outline on whole brain. **(B-D)** Confocal images of MB (magenta) and putative *Pdfr* (green) expression patterns. **(B)** Maximum intensity projection (MIP) of calyx, with single planar slice shown in **(C)** where dotted outline shows the only cell with colocalization of markers seen in n=3 brains (magnified, inset). **(D)** MIP of MB lobe region. **(E)** LNv neurons were activated by *pdf-GAL4* expression of the ATP-stimulated P2×2 channel and compared to AHL vehicle and empty driver controls. Data shows MB calyx intracellular cAMP as reported by EPAC1camps FRET signal imaged at 2 Hz; line mean, shadow ± SEM. N of each genotype (shown in parentheses) pooled from five timepoints throughout the 24 h cycle. T-test comparisons (paired, unpaired) of mean terminal 10 s are non-significant.

**Figure 4.**
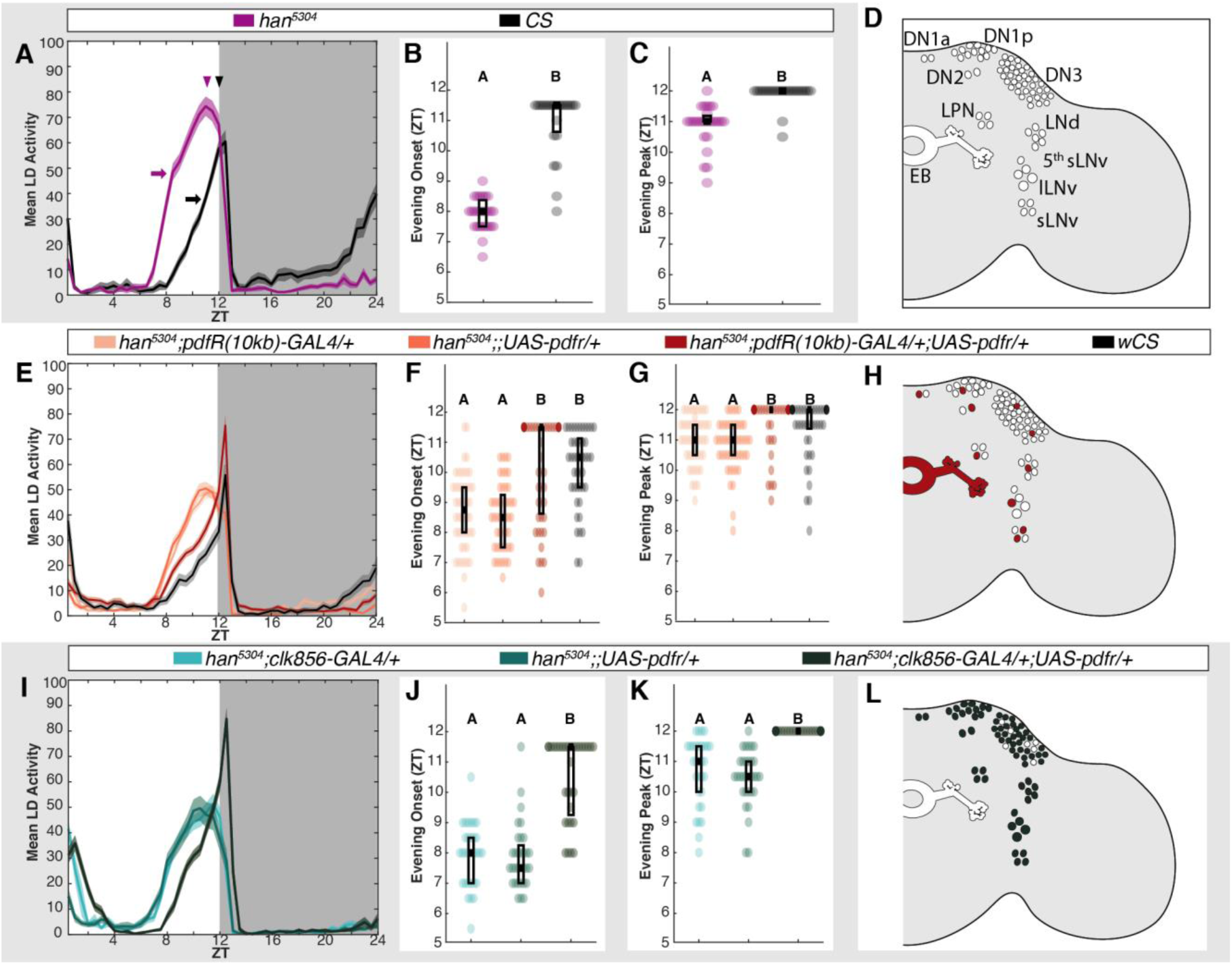
Clock-based PDFR supports normal locomotor activity. LD activity and appetitive STM for *han*^*5304*^ and cell-specific *Pdfr* rescue mutants. **(A**,**E**,**I)** Population LD activity (mean ± SEM) shown in 30 min bins averaged from 3 consecutive days. In (A), mean population values of evening anticipation onset (arrows) and evening peak (arrowheads) are shown. Evening anticipation onset **(B**,**F**,**J)** and the evening activity peak **(C**,**G**,**K)** were extracted for individual subjects (circle datapoints; overlaid box-and-whisker). **(D)** In addition to other cells, core clock neurons (categorized by subtype DN, LNd, LPN, and LNv) and EB soma and ring neuropil in *han*^*5304*^ lack PDFR (indicated by empty outlined ROIs). **(H**,**L)** Compared to (D), filled ROIs demonstrate (H) *PdfR(10kb)-GAL4* and (L) *clk856-GAL4* mediated PDFR rescue expression patterns on the *han*^*5304*^ background. Letter categorization indicates groups of statistical similarity (p > 0.05) or difference found by (B,C) Student’s unpaired t-test (p < 1E-7); (F,G,J,K) Kruskal-Wallis with Bonferroni posthoc comparisons (all significant comparisons panel F: p < 0.0001; panel G: p < 0.05; panel J: p < 1E-7; panel K p < 1E-9).

**Figure 5.**
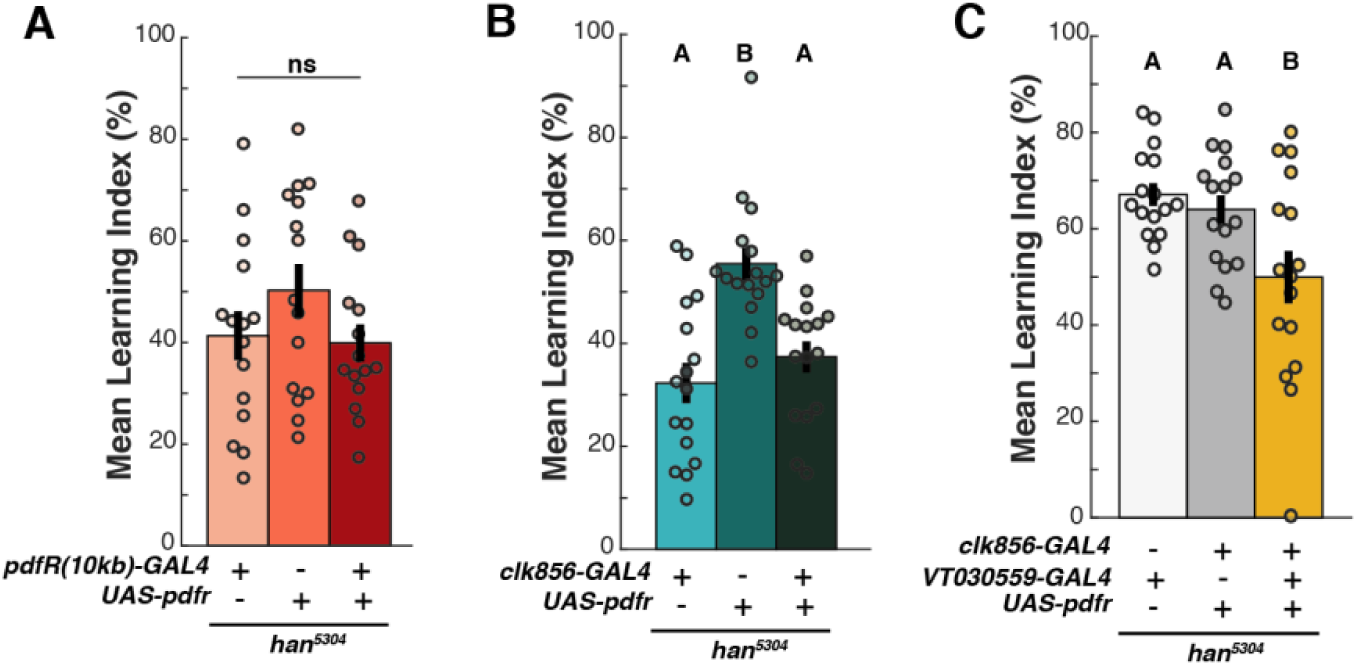
PDF signaling in the clock and MB does not support STM. Appetitive STM of *han*^*5304*^ mutants with cell-specific constitutive *Pdfr* expression. Mean LI scores are shown ± SEM, with individual datapoints (circles). Letter categorization within each panel indicates groups of statistical similarity (p > 0.05) or difference found by 1-way ANOVA with Bonferroni posthoc comparisons (all significant comparisons, panel B: p<0.005; C: p < 0.05).

**Figure 6.**
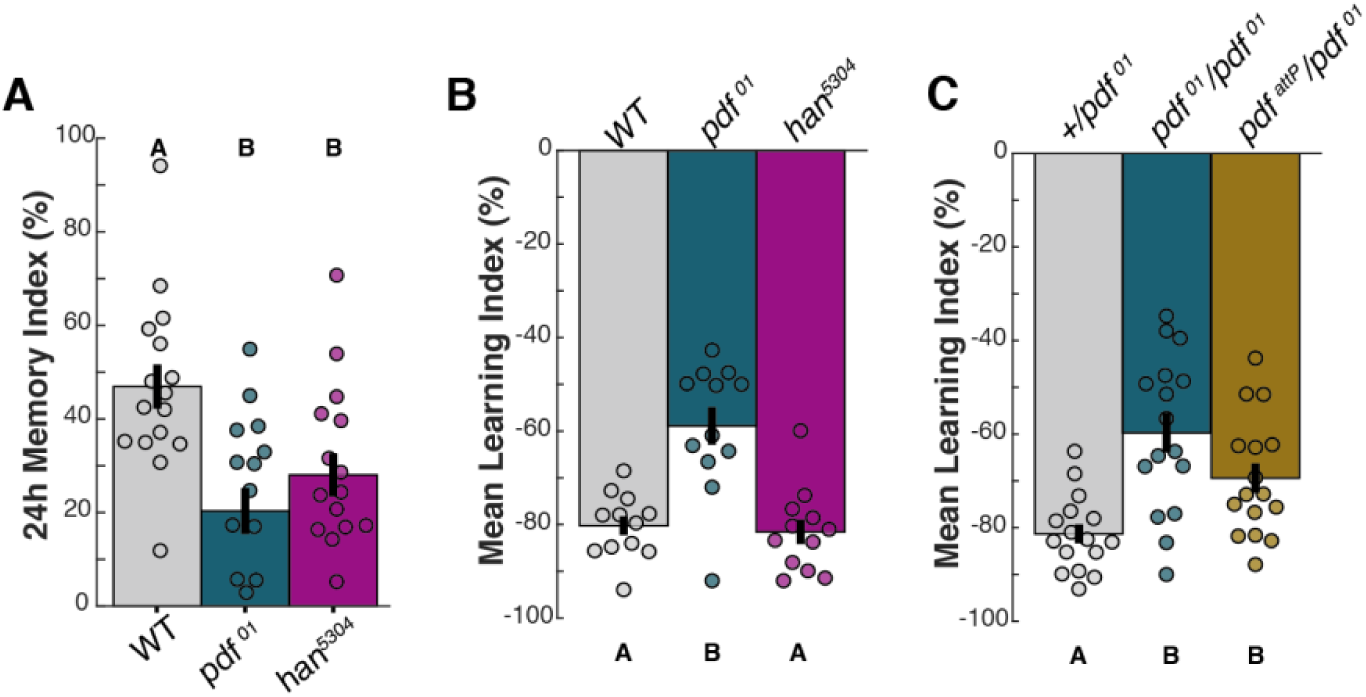
Multiple receptor targets permit valence-specific support of STM by a single clock output. Mean LI scores are shown ± SEM, with individual datapoints (circles). **(A)** 24 h appetitive LTM of *pdf*^*01*^ and *han*^*5304*^ mutants. **(B**,**C)** Aversive STM of WT and PDF pathway mutants. Letter categorizations indicate groups of statistical similarity (p>0.05) or difference found by 1-way ANOVA with Bonferroni posthoc comparisons (all significant comparisons, panel A: p<0.05; panel B: p<1E-4; panel C: p<0.05).

### Temporal panneuronal expression of Pdfr (elav-GS-GAL4, adult only)

GeneSwitch experimental flies (*han*^*5304*^;;*elav-GS-GAL4/UAS-Pdfr*) were housed after eclosion on normal yeast food containing 0.2g/L RU486 (Sigma, Cat#8046) diluted into 100% ethanol; control flies were housed after eclosion on normal food containing equal volume 100% ethanol. Total exposure time was 6-12d. Flies were starved for appetitive learning assays on nutrient-free agarose ± RU486 or ethanol control.

### Learning assays

Appetitive and aversive associative olfactory memory assays were performed in an environmental room in red light at 25°C with 65% ambient humidity. Flies were between 4-14 d old (STM) or 4-10 d old (LTM) and given at least 10 min acclimation to the environmental room prior to training or testing. Data for each experiment was pooled from at least three independent experimental days.

Appetitive learning assays were performed as previously described (Liu et al., 2012). Briefly, flies were starved to 10% mortality. Filter papers were prepared blank or with 2 M sucrose as null or unconditioned stimulus (US), and 10% MCH and OCT prepared as conditioned stimulus (CS) odors. 50-100 starved flies were loaded into a vial and exposed to a one trial training of sequential CS_A_-US and CS_B_-null pairings of 1 min (STM) or 2 min (LTM) duration. Flies were then either tested for CS preference (2 min STM) or placed into fresh food vials for 4 h and then starved for 20 h prior to CS preference testing (24 h LTM). Testing involved a 2 min exposure to both CS odors, after which flies choosing either odor were counted. A preference index (PI) was calculated for each trial as [(# of flies in CS_A_) - (# of flies in CS_B_)] / [(# of flies in CS_A_) + (# of flies in CS_B_)]. This PI was averaged with the PI of a temporally paired CS reciprocal trial to generate the final Learning Index (LI), calculated as a percentage. In this way, each datapoint shown for these experiments represents 100-200 flies.

Aversive learning STM assays were performed similarly to appetitive STM assays with the following modifications. Flies were not starved prior to training, and the US was provided by supplying 12 1-sec 90mV shocks during the 1 min CS-US pairing. As for appetitive STM, flies were immediately tested for CS preference and a final LI calculated from paired reciprocal tests.

### Activity Assays & Data Extraction

Male flies were collected immediately after eclosion and individually loaded into sleep tubes and detectors as in (Haynes et al., 2015). Flies were entrained for 3 d to 12:12 LD in 25°C incubators, after which 3 d of 12:12 LD baseline activity was collected followed by release into free-running DD conditions for 9 d. During this time locomotor activity data was collected using the Drosophila Activity Monitor (DAM; TriKinetics, Waltham MA) as previously described (Agosto et al., 2008). Subsequently, beam break counts were extracted using DAMFileScan (TriKinetics, Waltham MA) and analyzed for sleep and activity using MATLAB SCAMP (v.2019, Chris Vecsey, Skidmore) as done previously (Haynes et al., 2015). DD analysis was performed on days 2-8 DD data, with percent of population rhythmic values calculated using cutoff criteria where a rhythmicity index below 0.2 was considered arrhythmic. LD data was calculated from an average of days 1-3 in LD. Custom MATLAB scripts were written to extract evening anticipation onset and evening peak values from 30 m binned LD activity data of individual subjects and are available on GitHub (URL to be provided). Evening anticipation onset was calculated as the max value of the 3 bin rolling average of the first derivative for data between ZT6-11.5 (to prevent artifact from lights-off startle effects). Evening peak was calculated as the max value between ZT6-12.

### Imaging

Functional imaging (Fig 3E) was performed on *ex vivo* whole brain dissections with imaging hardware, solutions and acquisition software as described previously (Haynes et al., 2015). Planar image acquisition in MB calyx and surrounding soma was performed at 2Hz throughout a 3 min total experimental time for each brain, during which 30 s of baseline AHL perfusion was followed by 150s of ATP or AHL vehicle perfusion. Data was processed before analysis by custom MATLAB scripts which normalized each trace to its baseline and then fit and subtracted out any linear trends present in baseline due to bleaching (available on GitHub; URL to be provided). Expression pattern imaging (Fig. 3B-D, Multimedia 1,2) was performed as follows: Brains were fixed, stained, mounted and embedded according to Janelia FlyLight protocols (‘Dissection and Fixation 2% PFA’, ‘IHC-Double Label’, ‘DPX Mounting’; https://www.janelia.org/project-team/flylight/protocols). Primary antibodies included mouse monoclonal anti-GFP (1:200; Roche Applied Science, Cat#11814460001), and rabbit polyclonal anti-Ds-red (1:200; Clontech, Cat#632496). Secondary antibodies included Cy2 goat anti-mouse (1:400; Jackson, Cat#115-225-166) and AlexaFluor594 donkey anti-rabbit (1:400; Jackson, Cat#711-585-152) as recommended in Janelia protocols. Samples were imaged on a Zeiss LSM 880 microscope using a Plan-Apochromat 25x/0.8 Imm Corr DIC M27 objective. Data presented in Fig.3B,C and Multimedia Fig.1 were imaged using 561/595 and 488/515 channels with a slice depth of 0.65 μm and subsequently processed with Airyscan. Data presented in Fig.3D and Multimedia Fig.2 were acquired with a slice depth of 0.75 μm using 458/550 and 633/698 channels. 3D rendering was performed on Zeiss Zen 2.3 software.

### Experimental design and statistical analysis

All statistical analyses used are detailed in Results and Figure Legends and were performed using MATLAB R2019a (MathWorks, Inc.). As the Kolmogorov-Smirnov test for normality is known to be overly sensitive with small sample sizes, statistical approaches accommodating unequal variance or non-parametric data were used in cases (1) where maximum standard deviation (SD) was equal or greater than three times’ the minimum SD, or (2) there was a known ceiling/floor imposing non-normality. Within a figure, statistically similar groups (p>0.05) are identified by the same letter assignment; statistically different groups (p<0.05) can be identified by having differing letter group assignments. For all tests requiring posthoc comparisons (ANOVA, Kruskal-Wallis), the Bonferroni method was used. For any significant (p<0.05) posthoc comparison p-values of an analysis, the largest significant p-value informed the selection of the closest standard p-values ultimately reported in Figure Legends (i.e. p<1E-4 in the case where the largest p-value was 0.000045).

## RESULTS

### Wildtype appetitive olfactory STM is stable throughout the day

While it has previously been shown that there are time-of-day (TOD) effects on some forms of olfactory memory in *Drosophila* (Chouhan et al., 2015; Fropf et al., 2014, 2018; Lyons & Roman, 2009), there has been no investigation of the role of the clock in appetitive olfactory STM. To address this, we first tested appetitive STM of *Canton-Special* wildtype (WT) flies entrained to a 12:12 LD cycle at six timepoints evenly spaced throughout the 24 h day (Fig. 1A, black). Grouped flies (100-200) were exposed to two neutral odors, one paired with sucrose and the other unpaired. Memory of this experience was assessed directly after training by allowing animals to choose between the two odors; WT flies prefer the odor previously paired with sucrose (Tempel et al., 1983). While a double-plot of the mean learning index (LI) of each timepoint was best fit with a nonlinear 1-term Fourier curve (R^2^=0.9889; linear fit R^2^=0.5400), there were no statistically significant TOD changes (n=8 per timepoint, 2-way ANOVA, p_ZT_=0.0373; posthoc comparisons for WTxZT, all p=1.00). Thus, WT appetitive STM appears to be relatively stable throughout the 24 h circadian cycle.

### The magnitude and TOD-independence of appetitive STM requires PDF/PDFR signaling

To determine if the circadian clock has a role in regulation of appetitive STM, we asked if the major molecular output of the core clock, the neuropeptide Pigment-dispersing factor (PDF) (Renn et al., 1999; Shafer & Yao, 2014) and its receptor PDFR (Choi et al., 2009; Hyun et al., 2005; Lear et al., 2005; Mertens et al., 2005), affected this behavior. Mutants lacking PDF (*pdf*^*01*^) or PDFR (*han*^*5304*^) were tested for appetitive STM alongside WT flies (Fig. 1A). Both *pdf*^*01*^ and *han*^*5304*^ showed a clear STM deficit compared to WT at all time points (2-way ANOVA: p_genotype_ = 6.43E-23; Bonferroni posthoc comparisons: p_WT,pdf_ = 3.62E-23, p_WT,han_ = 4.31E-12, p_pdf,han_ = 3.44E-05). These animals had normal odor and sugar responses (data not shown). Given the requirement found for both PDF and its receptor PDFR, we conclude that the PDF signaling pathway supports STM throughout the day.

These *pdf*^*01*^ and *han*^*5304*^ data were also well fit by non-linear Fourier curves (R^2^ = 0.7812 and 0.8472 respectively; linear fit R^2^=0.0052 and 0.0987 respectively), but their apparent amplitudes of oscillation were larger than those of WT. This is made more obvious if the data are normalized to their means to allow direct comparison between genotypes (Fig. 1B). While this normalization damps the amplitude of the WT oscillation, the *pdf*^*01*^ and *han*^*5304*^ curves show even larger excursions, implying that the PDF signaling pathway may in fact stabilize STM formation over the course of the day.

To test this, we assayed WT STM compared to *pdf*^*01*^ or *han*^*5304*^ at the times of day where we had previously observed largest deviations from the mean: dawn (ZT1) and dusk (ZT13) (Fig. 1C,D). These experiments replicated the prior overall deficits seen in *pdf*^*01*^ and *han*^*5304*^ appetitive STM and confirmed that WT appetitive STM does not change with TOD (2-way ANOVA, posthoc comparisons: Fig. 1C p_WT1,13_=1.00; Fig. 1D p_WT1,13_ = 1.00). However, as predicted by our previous findings, both *pdf*^*01*^ and *han*^*5304*^ had significantly lower learning scores at ZT1 relative to ZT13 (2-way ANOVA, posthoc comparisons: Fig. 1C p_pdf1,13_ = 0.008; Fig. 1D p_han1,13_ = 0.003). The loss of PDF signaling through PDFR therefore uncovers strong TOD effects on the ability to learn. We conclude that PDF and its receptor PDFR are generally necessary for STM but also have a critical role in ensuring that animals can learn equally well at all times of day.

### Appetitive STM requires PDF signaling in adult neurons

In the mature adult fly, PDF is uniquely produced in the LNv neurons of the clock, but PDFR is present in both neurons and glia (Im & Taghert, 2010). To distinguish between a neuronal and a glial PDF target involved in STM, we asked if neuron-specific expression of a *Pdfr* cDNA would rescue the *han*^*5304*^ appetitive learning deficit. Panneuronal expression of *Pdfr* on a *han*^*5304*^ background using the *nsyb-GAL4* driver significantly increased STM compared to parental controls (Fig. 2A, 1-way ANOVA, p=4.61E-06; posthoc comparisons p_gal4,UAS_ = 0.808, p_gal4,UAS+gal4_ = 6.10E-06, p_UAS,UAS+gal4_ = 2.07E-04). We excluded the possibility of neomorphic effects from the broad *nsyb-GAL4* expression pattern by investigating the same panneuronal PDFR overexpression on a WT background, which failed to generate significant changes in STM compared to parental controls (data not shown). Therefore we concluded that PDF/PDFR signaling in neurons is sufficient for normal appetitive STM.

To determine if adult-specific PDF signaling was sufficient for normal STM, we used the drug-inducible driver *elav-GS-GAL4* to panneuronally express *Pdfr* on the *han*^*5304*^ mutant background solely during adulthood. Flies were placed onto food containing RU486 or EtOH vehicle directly after eclosion to fully induce transgene expression (Depetris-Chauvin et al., 2011; Osterwalder et al., 2001). Under these conditions, *han*^*5304*^;;*elav-GS>Pdfr* flies fed with RU486 (Fig. 2B) showed significantly increased appetitive STM compared to their vehicle-fed sibling controls and had learning scores equivalent to the concurrently run WT control (1-way ANOVA, p = 8.0235e-04; posthoc comparisons p_EtOH,RU486_ = 0.0144, p_EtOH,WT_ = 5.872e-04, p_RU486,WT_ = 0.3288). Taken together, we conclude that PDFR in a population of adult neurons promotes appetitive STM.

### Mushroom body Kenyon cells are not direct targets of PDF

We began our search for the relevant PDFR^+^ neurons in the mushroom body (MB), a site of integration essential for olfactory learning and memory in the fly. Olfactory information is delivered to the MB neuropil by sparse random activation of subpopulations of roughly 2000 Kenyon cells (KCs) whose soma surround the calyx of the MB. Within the compartmentalized MB neuropil lobes, KC axonal projections receive stimulus-specific information from dopaminergic inputs, allowing for time-space coincidence of odor-stimulus pairings (for review, see Busto et al., 2010). Dorsomedial projections from PDF^+^ sLNv clock cells skew posteriorly in close proximity to the MB KCs and calyx (Fig. 3A, and (Helfrich-Förster, 1995); while these projections contain small clear core vesicles (Yasuyama & Meinertzhagen, 2010) the recently published *Drosophila* adult brain connectome shows no synaptic routes between sLNv and MC KC neurons (Xu et al., 2020) permitting us to exclusively focus on sLNv extrasynaptic dense core vesicle signaling as a communication mechanism between sLNv and MBs. In fact, sLNv projections show TOD-dependent changes in PDF immunoreactivity (Park et al., 2000) and low levels of PDFR mRNA have been detected in some KC subpopulations (Crocker et al., 2016). These prior findings led us to ask if KCs could be a direct target of PDF.

We labeled KCs with membrane-bound RFP using the *VT030559-GAL4* driver and looked for colocalization with membrane-bound GFP expressed with *Pdfr-2A-LexA* (Deng et al., 2019), a gene-fusion in the *Pdfr* locus (Fig. 3B-D) which should recapitulate the endogenous expression pattern of *Pdfr*. Given the dense packing of the KC soma, we took particular care to maximize our confocal Z-resolution and visualize independent soma by utilizing AiryScan deconvolution. A Z-axis maximum image projection (MIP) of the KC soma and MB calyx (Fig. 3B) shows potential overlap of reporter signal; however a flythrough of the stacked dataset (Multimedia Fig. 1) shows that the entirety of this ‘overlap’ can be attributed to the extremely close proximity of the PDFR^+^ DN1 clock cell soma and PDFR^+^ DN1/LNv projections to the calyx and KC soma. In an examination of ∼6000 KCs from three independent brains we saw only a single cell that expressed both GFP and RFP (Fig. 3C, inset). We also examined the MB lobe region to detect any innervation of the neuropil by PDFR^+^ projections; again, though the Z-axis MIP implies potential overlap (Fig. 3D), examination of 3D rotational data (Multimedia Fig. 2) revealed no detectable PDFR^+^ signal within the MB lobe neuropil. GFP signal closest in proximity belonged to EB-projecting PDFR^+^ cells.

We also attempted to visualize a functional connection between the PDF^+^ LNvs and the MB. PDFR is a G-protein coupled receptor which activates Gαs to increase intracellular cAMP (Duvall & Taghert, 2012, 2013). To allow us to see PDF-dependent changes in cAMP we expressed the FRET-based reporter EPAC1cAMPs (Shafer et al., 2008) in the entirety of the MB with *MB247-LexA*. In these flies, exclusive activation of LNvs was accomplished by expressing ATP-gated P2X_2_ channels under the control of *pdf-GAL4* and perfusing dissected brains with ATP (Lima & Miesenböck, 2005; Yao et al., 2012). Though we performed experiments at five different timepoints spanning the 24 h circadian day, we failed to detect any significant response to ATP within the MB calyx or KC soma (pooled data, Fig. 3E; paired T-test, p = 0.054; unpaired T-test, p = 0.153) relative to vehicle or genetic controls. The PDFR^+^ neuron(s) involved in the regulation of appetitive STM must therefore be extrinsic to the MB, acting as interneurons downstream of the PDF^+^ LNvs to regulate the memory center.

### Intra-clock PDF signaling is not sufficient for appetitive STM

The two most extensively characterized functions of PDF are regulation of daily locomotor activity timing and maintenance of clock circuit intracellular cycling (manifest by rhythmic locomotor activity in constant conditions). Both of these functions are carried out by PDF activation of PDFR in neurons of the core clock. Because we found that PDF is required for STM but does not directly signal to memory-relevant MB KCs, we next considered PDF signaling within the clock itself as an indirect regulator of memory formation. If cycling of the core clock was required to produce an STM-promoting non-PDF output or if circadian-regulated locomotion itself was a factor in memory formation, intra-clock signaling might be memory-relevant. We therefore asked if PDFR expression sufficient for the rescue of canonical *han*^*5304*^ LD and DD locomotor phenotypes is also sufficient for STM rescue. To do this we first sought to identify clock-related GAL4 lines that were capable of rescuing these phenotypes, with later intent to examine their efficacy in the STM assay.

The *han*^*5304*^ LD locomotor activity phenotype has an advance in the timing of evening anticipation onset and an early evening peak activity (Hyun et al., 2005). We observed *han*^*5304*^ activity for 3 consecutive days in 12:12 LD using the Drosophila Activity Monitor (DAM) system and, consistent with prior reports, *han*^*5304*^ flies showed accelerated evening onset and evening peak activity compared to WT (Fig. 4A-C; Student’s unpaired T-test: Fig. 4B, p=8.45E-14; Fig. 4C, p=4.38E-08). We then rescued expression of *Pdfr* in *han*^*5304*^ using two different clock-related drivers: *PdfR(10kb)-GAL4* and *clk856-GAL4*, and recorded locomotor activity under similar conditions. The *PdfR(10kb)-GAL4* expression pattern (Fig. 4H) includes the ellipsoid body and 1-2 cells of each clock subset (Parisky et al., 2016). We found that *Pdfr* expression under the *PdfR(10kb)-GAL4* promotor was sufficient to delay the *han*^*5304*^ evening onset and return evening peak timing back to WT (Fig.4E-G) (Fig. 4F,G Kruskal-Wallis with Bonferroni posthoc comparisons, (F) p=5.56E-10, posthoc comparisons: p_gal4,UAS_ =1.00, p_gal4,UAS+gal4_ =5.44E-05, p_UAS,UAS+gal4_ =2.67E-06, p_gal4,WT_ =3.86E-05, p_UAS,WT_ =1.89E-06, p_UAS+gal4,WT_ =1.00; (G) p=9.60E-09, posthoc comparisons: p_gal4,UAS_ =1.00, p_gal4,UAS+gal4_ =2.86E-05, p_UAS,UAS+gal4_ =9.71E-08, p_gal4,WT_ =2.67E-02, p_UAS,WT_ =5.61E-04, p_UAS+gal4,WT_ =0.568). Similarly, clock-limited PDFR expression with *clk856-GAL4* (Fig. 4L) (Gummadova et al., 2009) significantly delayed the accelerated *han*^*5304*^ evening onset and evening peak timing (Fig. 4I-K) (Fig. 4J,K Kruskal Wallis with Bonferroni posthoc comparisons, (J) p=1.58E-10, posthoc comparisons: p_gal4,UAS_ =1.00, p_gal4,UAS+gal4_ =1.24E-07, p_UAS,UAS+gal4_ =3.40E-09; (K) 1.94E-13, posthoc comparisons: p_gal4,UAS_ =0.772, p_gal4,UAS+gal4_ =8.39E-09, p_UAS,UAS+gal4_ =2.96E-12).

The other phenotype of the *han*^*5304*^ mutant is an inability to maintain circadian rhythmicity of locomotor activity after transfer to total darkness (Hyun et al., 2005). Although our STM assays utilize flies entrained to a 12:12 LD cycle, it was still possible that a PDFR^+^ component of the clock involved in maintenance of circadian rhythmicity was required for STM. We therefore examined the free-running activity of those flies in DD conditions, analyzing activity from DD days 2-8 (Table 1). Compared to WT, *han*^*5304*^ had significantly decreased rhythmicity index (RI) (Student’s unpaired T-test, p=2.84E-08) and periodicity (Welch’s T-test, p=0.0363) and the overall population was much more arrhythmic (70.4 % vs 95.6 %). PDFR expression with the *pdfR(10kb)* promotor failed to increase *han*^*5304*^ RI (1-way ANOVA, p=4.75E-14, posthoc comparisons p_gal4,UAS_ =0.19, p_gal4,UAS+gal4_ =0.0280, p_UAS,UAS+gal4_ =1, p_gal4,WT_ =4.38E-14, p_UAS,WT_ =1.05E-08, p_UAS+gal4,WT_ =5.575E-07), periodicity (Kruskal-Wallis, p=2.31E-05, posthoc comparisons p_gal4,UAS_ =1, p_gal4,UAS+gal4_ =1, p_UAS,UAS+gal4_ =1, p_gal4,WT_ =1.79E-04, p_UAS,WT_ =9.60E-05, p_UAS+gal4,WT_ =6.23E-03) or percent rhythmic values to WT levels. *han*^*5304*^;*clk856*>*Pdfr* flies had significantly increased RI (1-way ANOVA, p=1.61E-08, posthoc comparisons p_gal4,UAS_ =0.013, p_gal4,UAS+gal4_ =8.37E-04, p_UAS,UAS+gal4_=8.16E-09) and percent rhythmic population values compared to their parental controls, while periodicity showed decreased variance but no rescue in mean values (Kruskal-Wallis, p=1.49E-03, posthoc comparisons p_gal4,UAS_ =0.263, p_gal4,UAS+gal4_ =9.27E-04, p_UAS,UAS+gal4_=0.219).

**Table 1.**
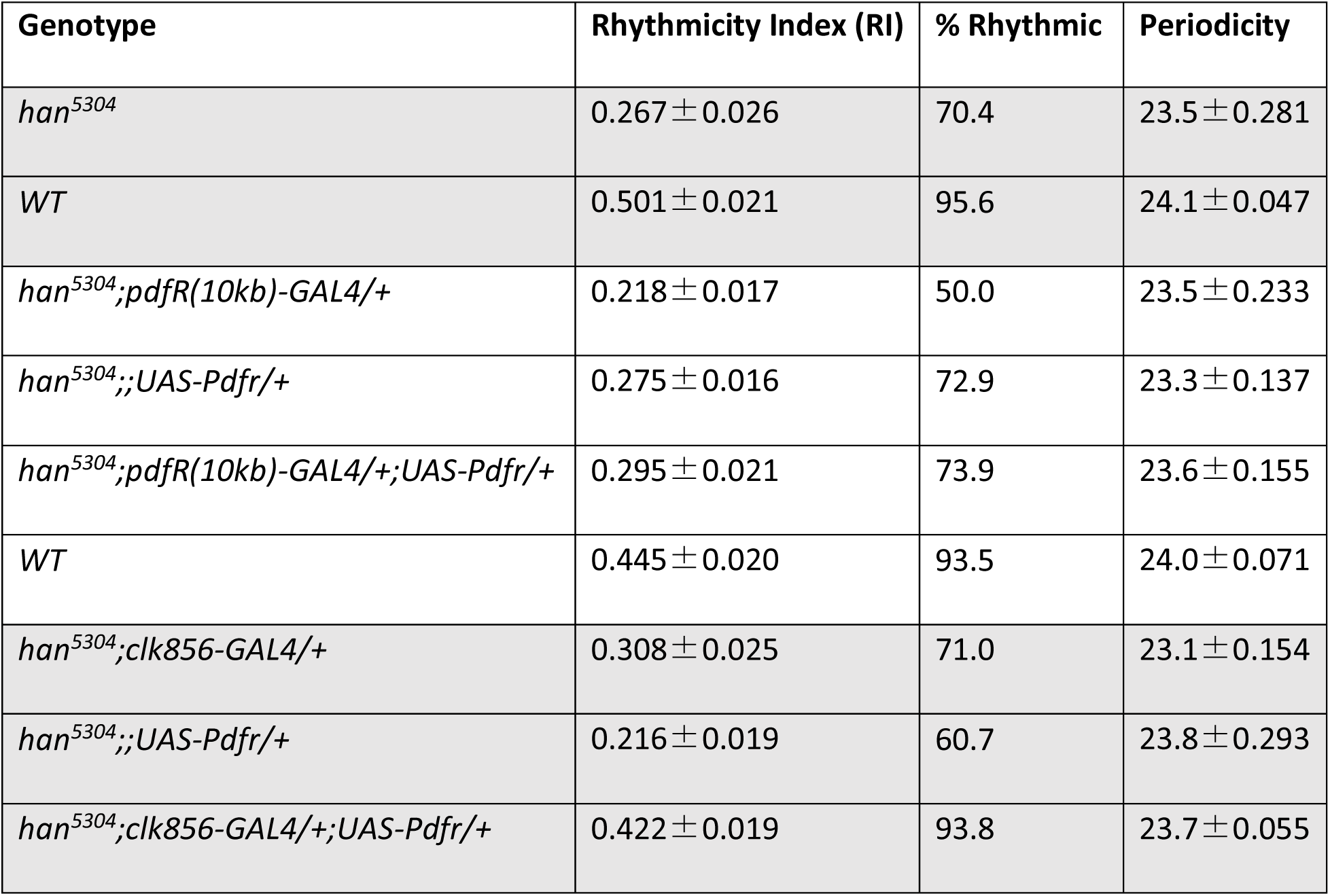
Free-running circadian locomotor activity properties of *han*^*5304*^ mutants and rescue genotypes. Flies used in Figure 4 were released into free-running DD conditions for 8 days following Figure 4 LD recordings; circadian properties are calculated from DD days 2-8. Shaded groupings indicate independent experimental comparison groups.

Taken together, we find that *pdfR(10kb)-GAL4* driven expression of PDFR rescues only LD activity phenotypes of *han*^*5304*^ mutants, while *clk856-GAL4* PDFR expression is sufficient to rescue both LD and DD phenotypes. These drivers therefore allowed us to ask whether PDF’s role in appetitive STM is mediated through core-clock-driven LD locomotor patterns or maintenance of circadian rhythmicity. Both of these hypotheses were disproven as we found that neither *pdfR(10kb)-GAL4* nor *clk856-GAL4* expression of PDFR was able to rescue the *han*^*5304*^ appetitive STM deficit (Fig. 5A,B; 1-way ANOVA, (A) p = 0.239, (B) p = 3.21E-05, posthoc comparisons: p_gal4,UAS_ =4.19E-05, p_gal4,UAS+gal4_ =0.879, p_UAS,UAS+gal4_ =0.001). Furthermore, as the *clk856-GAL4* expression pattern includes nearly all core clock neurons, it’s reasonable to entirely infer that clock-based PDFR expression alone is insufficient to support normal appetitive STM. However to fully exclude any possible synergistic interaction of PDF with the clock and MB in STM behavior, we co-expressed PDFR in the clock and MB simultaneously using *clk856-GAL4* and the strong KC-specific driver *VT030559-GAL4* on a *han*^*5304*^ background (Fig. 5C). This manipulation not only failed to rescue appetitive STM compared to parental controls, but in fact further impaired learning, likely due to either genetic background effects or to a neomorphic effect of strong overexpression of PDFR in the MB (1-way ANOVA, p = 6.15E-03, posthoc comparisons: p_VTgal4,UAS_ =1.00, p_VTgal4,clkgal4UAS+VTgal4_ =8.17E-03, p_clkgal4UAS,clkgal4UAS+VTgal4_ =0.0379). These data imply that PDF must be acting on a PDFR^+^ neuronal target exclusive from both the core clock and the MB KCs to regulate appetitive STM. Our results also argue that the multiple behavioral roles of PDF signaling (on locomotor activity, DD rhthyms, and memory) are independently controlled by the expression of PDFR in distinct PDF target neurons.

### Aversive olfactory STM requires PDF but not PDFR

Differential distribution of PDFR, the only known receptor for PDF, within and external to the clock allow it to have many different and independent behavioral roles. Other signaling pathways, however, often use an additional strategy for diversification: multiple receptors. We were struck by the difference in STM scores between *pdf*^*01*^ and *han*^*5304*^ mutants (Fig. 1A)—the *pdf*^*01*^ STM phenotype is significantly and consistently more severe than *han*^*5304*^. Since both of these alleles are protein null (Hyun et al., 2005; Renn et al., 1999) in theory they should have the same magnitude of deficit if PDFR is the sole receptor for PDF. If, however, there were a second receptor for PDF, it could provide some PDF signaling in the *han*^*5304*^ mutant and result in a milder phenotype.

To further explore the learning and memory roles of PDF and PDFR and determine if the disparity in STM deficits was common to other types of memory, we assayed appetitive long-term memory (LTM) and aversive STM. We first investigated a possible difference between *pdf*^*01*^ and *han*^*5304*^ 24 h appetitive LTM (Fig. 6A). While *pdf*^*01*^ flies were slightly more impaired than *han*^*5304*^ flies were, there was no statistically significant disparity between the mutants (1-way ANOVA, p = 8.00E-04, posthoc comparisons: p_WT,pdf01_= 7.79E-04, p_WT,han5304_ = 0.0215, p_pdf01,han5304_ = 0.770). However we found strong evidence for the existence of a second PDF receptor when we tested *pdf*^*01*^ and *han*^*5304*^ mutants for aversive STM (Fig. 6B). In this assay, flies are tested for memory to an odor that was previously paired with shock; instead of approaching the paired odor (as in the appetitive assay), flies demonstrate memory by avoiding the paired odor. As before, *pdf*^*01*^ mutants had impaired aversive STM compared to WT, but quite surprisingly, *han*^*5304*^ mutants showed no aversive STM deficit at all (1-way ANOVA, p = 5.08E-06, posthoc comparisons: p_WT,pdf01_= 4.90E-05, p_WT,han5304_ = 1.00, p_pdf01,han5304_ = 1.97E-05). To definitively confirm a requirement for PDF in aversive STM and rule out the influence of genetic background in *pdf*^*01*^, we crossed *pdf*^*01*^ flies to WT, *pdf*^*01*^, and the recently created *pdf*^*attP*^ mutant (Deng et al., 2019) and tested progeny for aversive STM (Fig. 6C). *pdf*^*01*^/+ heterozygotes were indistinguishable from WT for STM, confirming the recessive nature of the defect. *pdf*^*01*^ homozygotes and *pdf*^*attP*^/*pdf*^*01*^ transheterozygotes had significant STM defects (Kruskal-Wallis, p =4.78E-04, posthoc comparisons: p_CS/pdf01,pdfattP/pdf01_ = 0.0269, p_CS/pdf01,pdf01/pdf01_ = 3.91E-04, p_pdfattP/pdf01,pdf01/pdf01_ = 0.676). Collectively, these data suggest that PDF normally promotes aversive STM by acting *not* on PDFR, but on a novel, and still unidentified, receptor.

## DISCUSSION

A role for the circadian clock in cognition and memory has been demonstrated in many ways and in many organisms, including humans (for review, see Gerstner & Yin, 2010; Krishnan & Lyons, 2015; Smarr et al., 2014). To date, efforts to understand these phenomena have largely involved manipulations of intracellular molecular oscillator components such as the *period (per)* gene to abolish intracellular clock cycling. However this type of manipulation is not fully representative of the nature of human circadian misalignment disorders, as a role for intercellular signaling has also been shown (Yamaguchi et al., 2013). In *Drosophila*, where investigators have tools to interrogate highly conserved circadian processes in great mechanistic detail, manipulation of the signaling peptide PDF allowed us to specifically examine the role of a key neuromodulatory output of the core clock circuit in cognition. Our work demonstrates a requirement for PDF in the formation of associative olfactory memory (schematized in Figure 7). Putting our data in the context of what is known about the molecular clock’s influence on cognition, we explore below the importance of TOD on WT and PDF mutant STM and provide evidence which supports the existence of a second PDF receptor.

**Figure 7.**
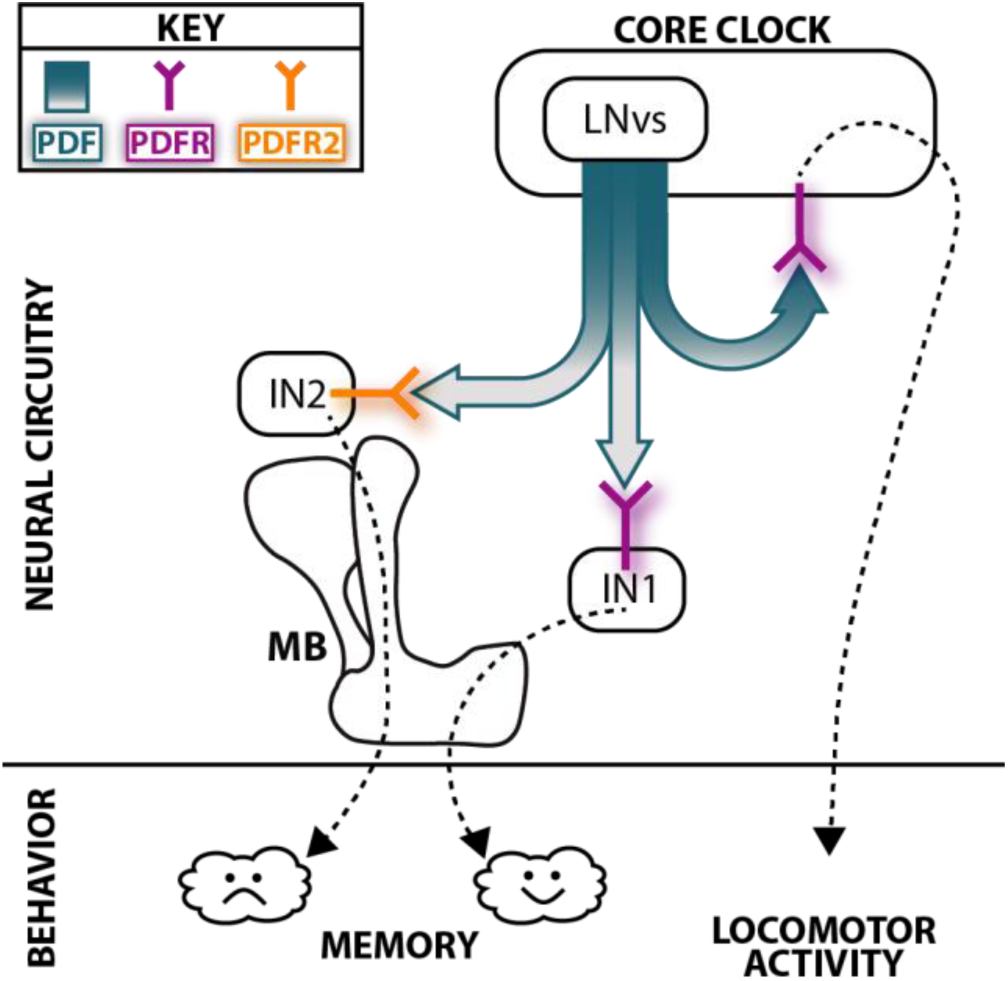
The core clock regulates distinct behaviors via discrete PDF targets: a proposed model. Localization of PDFR to the clock permits control of locomotor activity independently from control of memory. Signaling through PDFR in a population of interneurons extrinsic to both the clock and MB (here shown as IN1) permits regulation of appetitive memory. We propose that, in place of PDF-PDFR signaling, PDF activation of a novel unidentified receptor (here called PDFR2) in a separate population of interneurons (IN2) is required for aversive memory.

### PDF acts on multiple targets to regulate different behaviors

In the adult fly, we and others have shown that PDF released from the LNv neurons of the core clock circuit acts, via PDFR, both on the clock circuit itself and outside of the clock to drive key circadian features and rhythmicity of locomotor activity (Fig. 4, Table 1). Our data show that PDF and PDFR are also required for robust formation of appetitive olfactory memory and for equalizing the ability to form memory across the day (Figs. 1,2). Surprisingly, the intra-clock circuit PDF signaling pathway, which is sufficient for normal locomotor activity, plays no role in PDF’s regulation of memory, demonstrating that PDF independently directs multiple behaviors depending on the location of its target receptor PDFR. Regulation of associative olfactory memory requires PDF signaling outside both the clock and the MB KCs, a key memory formation center (Figs. 3-5). An additional layer of valence-specific control over memory formation is provided by the fact that PDF appears to act in aversive olfactory memory formation independently of PDFR, implying the existence of a second receptor system (Fig. 6).

### The role of core-clock driven circadian rhythms themselves in olfactory STM formation is limited

In *Drosophila*, a requirement for molecular clock components like *per* in cognitive tasks has varied, dependent on the type of learning and stage of memory studied (Chouhan et al., 2015; Fropf et al., 2014, 2018; Gailey et al., 1991; Inami et al., 2020; Le Glou et al., 2012; Lyons & Roman, 2009; Sakai et al., 2004). Interestingly, in cases where the locus of action for *per* has been mapped, PER acts outside the core clock circuit to regulate memory but these studies do not explore TOD effects. In studies which examine TOD effects on cognition (Fropf et al., 2014, 2018; Lyons & Roman, 2009), *per* mutants or animals rendered arrhythmic by altered light conditions lose TOD differences. Thus the role of the cycling core clock itself in learning and memory has been mostly restricted to TOD modulation, and PER, like PDF and PDFR, appears to have a role in memory that is independent of its function in the core clock circuit.

### PDF signaling opposes a latent TOD rhythm in appetitive STM

Our data suggest that the requirement for PDF in supporting normal levels of appetitive STM is TOD-independent i.e. PDF is required at all times of day to form robust appetitive STM. PDF has been implicated in a wide variety of behaviors that are influenced by circadian time (Chen et al., 1999; Chen et al., 2016; Chung et al., 2009; Keene et al., 2011; Kim et al., 2013; Krupp et al., 2013; Mertens et al., 2005; Nagy et al., 2019) but its only previously known roles in learning and memory are courtship-related (Inami et al., 2020). The PDF peptide is anatomically well placed to be an agent of general learning enhancement since it can act on long range targets by virtue of extrasynaptic release and diffusion, allowing it to provide neuromodulation over the entire circadian cycle. The fact that PDF has been characterized as an arousal-promoting signal (Chung et al., 2009; Parisky et al., 2008) may be germane to this general role since arousal state is critical to attention state in working memory (Ricker et al., 2018).

An intriguing additional feature of the *pdf* and *Pdfr* mutant phenotypes is the emergence of a latent TOD oscillation in STM (Fig. 1). Daily changes in sLNv structure and cycling accumulation of PDF peptide at sLNv terminals (Park et al., 2000) have minima and maxima at ZT1 and ZT13 (Fernández et al., 2008; Gorostiza et al., 2014) which align with our observed peak and trough of appetitive STM in PDF signaling mutants. The role of PDF signaling in morning arousal (Parisky et al., 2008) may explain the early day appetitive STM trough of *pdf* and *Pdfr* animals: since a heightened arousal state facilitates learning, an absence of PDF signaling reduces morning arousal and cripples the ability of animals to make associations with food-predictive odors. Furthermore, the latent nature of TOD effects in PDF signaling mutants relative to the stable nature of WT STM throughout the day allows us to infer that this stability is the sum of oscillating contributions from PDF signaling and one or more other cycling factors. This unknown oscillator may be complex, dependent on interactions between sleep drive, the sleep homeostat and arousal systems. In humans, multiple oscillators for working memory have been proposed (Folkard et al., 1983).

### TOD-independent appetitive memory has survival value

The diversity and abundance of roles for PDF in behavior illustrates how essential the output of the core clock is to an organism’s optimal function and survival. It’s important nevertheless to explore why, in the context of appetitive STM (which is ultimately TOD-independent), an organism’s capacity for appetitive learning should be so critically linked to its timekeeping capabilities. Though prior work is conflicting regarding TOD influence on associative STM (Fropf et al., 2018; Lyons & Roman, 2009), TOD effects are consistently found in associative LTM (Chouhan et al., 2015, 2017; Fropf et al., 2014, 2018; Lyons & Roman, 2009). In this context we ask: why is it important for appetitive learning to remain stable throughout the day? Given the nutrient-based nature of our assay, we posit that at the time of training, an organism lacks information necessary to evaluate the benefit of devoting metabolic resources to consolidation of that learning—i.e. future food availability is unknown. Though the widely-used appetitive STM paradigm requires starvation for expression of memory immediately after training, significant expression of appetitive LTM requires starvation only at the time of retrieval but not at the time of acquisition (Chouhan et al., 2017; Krashes et al., 2009). Furthermore, feeding after appetitive training prompts the decay of appetitive STM within 10-30 min, though LTM can still be observed with subsequent starvation (Krashes et al., 2009). In light of a STM requirement for PDF throughout the day, we hypothesize that PDF may help maximize information acquisition at the front end of the process, to allow memory to be dispensed with or consolidated in a manner dependent on future internal metabolic state information.

### PDF signaling in aversive memory utilizes an unknown receptor

We also find that PDF is required for aversive olfactory STM. Surprisingly, however we did not uncover a requirement for PDFR, the only characterized receptor for this peptide. Aversive associative STM is therefore the only behavior known for which PDF is required but PDFR is dispensable, providing the first evidence in *Drosophila* for a second PDF receptor. In other organisms, circadian output peptides have multiple receptors. Three GPCRs for the *C. elegans* homolog of PDF have been identified, each of which is highly similar to *Drosophila* PDFR and related to VIPR1 and VIPR2, the two known mammalian receptors for VIP (a mammalian peptide with similar clock roles) (Janssen et al., 2008, 2009). We suggest that receptor diversity allows the valence-specific regulation of associative STM by regionally segregated and distinct receptors. Since the circuitry involved in acquisition of appetitive versus aversive memory involves different sets of neurons (Liu et al., 2012; Riemensperger et al., 2005; Yamazaki et al., 2018), this implies that that PDF will act at circuit nodes in each pathway that are valence-specific and is consistent with our finding that PDF does not act on Kenyon cells, the final common substrate of memory. Ultimately, identification and characterization of the second PDF receptor will be required to fully understand this peptide’s multiple roles in behavior.

The authors declare no competing financial interests.

## ACKNOWLEDGEMENTS

This work was supported by NIH R01 MH067284 to LCG, NIH F31 NS110273 to EJRR and NIH T32 NS007292. We offer our thanks to Dr. Timothy D. Wiggin for his original versions of EPAC data analysis scripts and his excellent feedback on the project overall.

## FIGURE LEGENDS

**Multimedia Figure 1. Putative PDFR expression is excluded from the MB Kenyon cells.** Confocal image of MB calyx and Kenyon cell soma, acquired with a slice depth of 0.65 μm, processed with Airyscan, and rendered as a fly through 5 fps video (posterior → anterior). MB calyx and Kenyon cell soma are shown in magenta (fixed with dsRed antibody staining of *VT030559-GAL4>UAS-mCD8-IVS-RFP*); putative PDFR expression pattern is shown in green (fixed with GFP antibody staining of *pdfR-2A-LexA>LexAop-mCD8-IVS-GFP*).

**Multimedia Figure 2. Putative PDFR expression is excluded from the MB lobe neuropil.** Confocal image of MB lobe neuropil, acquired with a slice depth of 0.75 μm and rendered as a 3D rotation around the X axis. MB lobes are shown in magenta (fixed with dsRed antibody staining of *VT030559-GAL4>UAS-mCD8-IVS-RFP*); putative PDFR expression is shown in green (fixed with GFP antibody staining of *pdfR-2A-LexA>LexAop-mCD8-IVS-GFP*). The MB β/β’ lobe tips hug the PDFR^+^ EB projection rings without overlap.

